# Lipidomic Analysis Reveals Drug-Induced Lipoxins in Glaucoma Treatment

**DOI:** 10.1101/2025.01.24.634771

**Authors:** DJ Mathew, S Maurya, J Ho, I Livne-Bar, D Chan, Y Buys, M Sit, G Trope, JG Flanagan, K Gronert, JM Sivak

## Abstract

Synthetic prostaglandin analogues, such as latanoprost, are first-line treatments to reduce intraocular pressure (IOP) in the management of glaucoma, treating millions of patients daily. Glaucoma is a leading cause of blindness, characterized by progressive optic neuropathy, with elevated IOP being the sole modifiable risk factor. Despite this importance, the underlying latanoprost mechanism is still not well defined, being associated with both acute and long term activities, and ocular side effects. Prostaglandins are eicosanoid lipid mediators. Yet, there has not been a comprehensive assessment of small lipid mediators in glaucomatous eyes. Here we performed a lipidomic screen of aqueous humour sampled from glaucoma patients or healthy control eyes. The resulting signature was surprisingly focused on significantly elevated levels of arachidonic acid (AA) and the potent proresolving mediator, lipoxin A_4_ (LXA_4_) in glaucoma eyes. Subsequent experiments revealed that this response is due to latanoprost actions, rather than a consequence of elevated IOP. We demonstrated that increased LXA_4_ inhibits pro-inflammatory cues and promotes TGF-β_3_ mediated tissue remodeling in the anterior chamber. In concert, an autocrine prostaglandin circuit mediates rapid IOP-lowering. This work reveals parallel mechanisms underlying acute and long-term latanoprost activities during the treatment of glaucoma.

## INTRODUCTION

Glaucoma represents a spectrum of diseases characterized by progressive retinal ganglion cell degeneration, optic neuropathy and visual field loss, with elevated intraocular pressure (IOP) being the sole modifiable risk factor (1, 2). Glaucoma is a leading cause of irreversible blindness and is estimated to affect over 110 million people by 2040 (3). Reduction of IOP is the standard of care in glaucoma treatment and is achieved using medical, laser, or surgical management. These approaches either increase the outflow of aqueous fluid from the anterior chamber of the eye or reduce fluid production. However, the relationship between IOP and glaucoma progression is complex. For example, a lowered mean IOP is not always a reliable indicator of disease stability, and increased risk of glaucoma progression is associated with a higher diurnal variation in IOP (4). In addition, the risk of disease progression is higher for the same IOP in more advanced stages of glaucoma (5). Therefore, it is critical to uncover additional biochemical mediators driving glaucoma pathogenesis and their links to IOP regulation.

The most commonly prescribed classes of IOP-lowering medications are prostaglandin analogues, followed by beta adrenergic blockers, carbonic anhydrase inhibitors and alpha-2 adrenergic agonists. Prostaglandin analogues (PGAs), such as Latanoprost, Bimatoprost and Travoprost are the most commonly used first-line agents for medical management (6–8), treating millions of glaucoma patients daily (9). These drugs increase both trabecular and uveoscleral aqueous humor outflow (10, 11) through two proposed mechanisms; FP receptor-mediated ciliary muscle relaxation (12), and also through increased permeability of outflow tissues via TGF-β mediated matrix metalloproteinase (MMP) activity (13–16). Despite their widespread use, the detailed biochemical mechanisms linking PGA actions to these dual IOP effects remain poorly understood, particularly the explanation behind recently reported long-term IOP-lowering after cessation of treatment (14, 17, 18). Although generally well-tolerated, long term PGA treatment is accompanied by diverse and well-documented ocular and periocular adverse side effects, such as changes to eyelid and iris pigmentation, hyperemia, iritis, corneal thinning, eyelash growth, periorbital fat atrophy, and potential associations with macular edema and uveitis (19–21). Therefore, it has become increasingly important to unravel the underlying drug mechanisms in order to potentially uncouple and target these activities separately.

Endogenous prostaglandins are lipid mediators with key roles in normal physiology and drive or amplify inflammatory responses. They are enzymatically generated from arachidonic acid (AA) by cyclooxygenases (COX-1 and COX-2) (22). Prostaglandins are part of an intrinsic eicosanoid (AA metabolite) network in tissues that also include bioactive lipoxygenase (LOX) metabolites, such as the lipoxins that are produced by the actions of 5-LOX and 12/15-LOX (23). In constrast to prostaglandins, the lipoxins A_4_ and B_4_ (LXA_4_ and LXB_4_) are themselves potent mediators of inflammation resolution and cellular homeostasis (24–26). LXA_4_ signaling and production has been linked to a variety of ocular surface and inflammatory diseases (26, 27), and we recently demonstrated that therapeutic lipoxin supplementation resulted in structural and functional neuronal rescue in rodent glaucoma models (28–30), and reduced neuroinflammation (26, 31, 32). Interestingly, formation of lipoxins, or related specialized proresolving mediators (SPM), can also be triggered pharmacologically, for example by statins or aspirin acetylation of COX-2 (33–38). Surprisingly, despite these central roles, the production of these and other small lipid mediators have not been well studied in glaucoma patients.

Given this background of clinical pharmacology, shared substrates and potential interactions between prostaglandins, lipoxins, and other lipid mediator circuits, we decided to profile these signals for the first time in glaucoma patients. Here we present a metabolomic characterization of LOX-and COX-generated mediators in aqueous humour sampled from patients with glaucoma compared to healthy controls. Unexpectedly, the resulting glaucoma patient signature was tightly focused on significantly elevated levels of AA and LXA_4_. Our subsequent experiments investigated the regulation of this AA-LXA_4_ circuit to reveal novel insights into the latanoprost mechanism of action.

## MATERIALS AND METHODS

### Patient recruitment and sample collection

Glaucoma patients with a diagnosis of primary open angle glaucoma (POAG), aged 60-80 years, and scheduled for glaucoma surgery with or without cataract surgery at Toronto Western Hospital or Kensington Eye Institute were approached for inclusion in the study. Age-matched control samples were obtained from patients without glaucoma undergoing routine cataract surgery. Patients with diabetes mellitus, systemic inflammatory disease, uveitis, retinopathy and age-related macular degeneration, or those taking non-steroidal anti-inflammatory drugs were excluded. From each eye, 100 μL of aqueous humor was collected using a 30 Gauge needle mounted on a 1-mL syringe, introduced into the anterior chamber anterior to the limbus, prior to any surgical intraocular entry. The samples were immediately snap frozen on dry ice and stored at -80°C until assessment by lipidomic analyses. All participants signed an informed consent form. This study was performed according to a protocol approved by the Research and Ethics Boards of University Health Network and Kensington Eye Institute and adhered to the tenets of the Declaration of Helsinki.

### Lipidomic Analyses

The lipid mediator profiles of collected aqueous humor, cell media, or rat tissue samples were analyzed by liquid chromatography (Agilent 1200 Series HPLC)-mass spectrometry (LC-MS/MS, QTRAP 4500, AB Sciex). The analyses included polyunsaturated fatty acids (AA, DHA and EPA), their downstream mediators (prostaglandins, leukotrienes, lipoxins, resolvins and maresins) and their metabolic precursors (monohydroxy-PUFA) and metabolites as previously published (28, 39–41). Note; these analytes are structurally and functionally distinct from membrane phospholipids, assessed in a single previous study of glaucomatous aqueous humor (42). Deuterated internal standards (PGE_2_-d4, LTB_4_-d4, 15-HETE-d8, LXA_4_-d5, DHA-d5, and AA-d8) were added to all samples before processing to calculate class-specific recoveries. Tissues were homogenized in a refrigerated bead homogenizer. Supernatants were extracted using C18 solid-phase columns. MS analyses was carried out in negative ion mode, and PUFA and their metabolites were quantitated by scheduled multiple reaction monitoring (MRM) using 3 to 4 specific transition ions for each analyte with a signal-to-noise ratio for the signature ion above 5:1 for raw MRM chromatograms. Quantification, calibration curves and HPLC retention times for each analyte were established with authentic synthetic standards (Cayman Chemicals) (Supplementary Figure 1).

### Human trabecular meshwork cell culture

Human TM-1 cells were cultured in low glucose Dulbecco’s Modified Eagle Medium (Sigma #D-5523), 10% premium fetal bovine serum, 2mM L-glutamine, 50 μg/mL gentamicin sulfate and Primocin (Invivogen category code ant-pm-1). We gratefully acknowledge Dr. Donna M. Peters for providing us with these well-characterized cells (43). Cells were grown in 10 cm plates at 8% CO_2_ and media was changed every two days. Upon reaching confluence, cells were treated with latanoprost (CAS No. 130209-82-4, Millipore Sigma), timolol maleate (CAS No. 26921-17-5, Millipore Sigma), dorzolamide (CAS No. 120279-96-1, Millipore Sigma) or brimonidine (CAS No. 59803-98-4, Millipore Sigma) for one hour at the indicated concentration. These drugs were dissolved in DMSO to the following concentrations: latanoprost, 0.5, 5, and 50 μM; timolol, 1, 10 and 100 μM; dorzolamide 0.5, 5, 50 μM; brimonidine 2, 20, 200 μM. Following treatment cells were collected after one hour for quantitative polymerase chain reaction (qPCR) and the cell culture media was collected and snap frozen for lipidomic analyses.

### Reverse Transcription-Quantitative Polymerase Chain Reaction (RT-qPCR)

RNA was extracted from TM-1 cells using the RNEasy Mini Kit (Qiagen, Cat. No. 74104) according to the manufacturer’s instructions. RNA samples were treated with RNase-free DNase (Promega RQ1 kit, Cat. No. PR-M6101). RNA purity was assessed using Nanodrop 2000 spectrophotometer (ThermoFisher Scientific, Cat. No. ND-2000), followed by cDNA synthesis using SuperScript IV First-Strand Synthesis System (Invitrogen Cat. No. 18091050). RT-qPCR was performed using SYBR-Green PCR Master Mix (Applied Biosystems, ThermoFisher Scientific, Cat. No. 4309155) on Eppendorf Realplex_2_ Mastercycler. The genes and primers used are listed in Table 1. Amplification of mRNA was normalized to GAPDH and the 2^-ΔΔCt^ comparative quantification method was used (44).

**Table 1:**
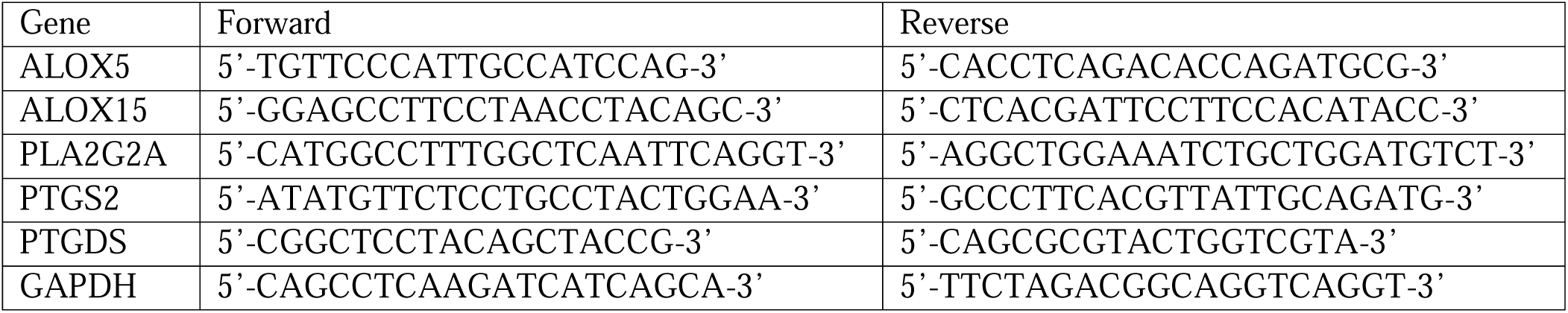
Primers used for Reverse Transcription-Quantitative Polymerase Chain Reaction (RT-qPCR)

### Animal Experiments

All procedures and protocols conformed to the guidelines of the ARVO statement for the use of animals in ophthalmic and vision research, and were approved by the University Health Network Animal Care Committee. All procedures were performed in accordance with all relevant regulations and are reported in accordance with ARRIVE guidelines. For all rodent experiments, six-week-old Long Evans rats (Charles River Laboratories, Massachusetts, USA) were used. The gradual ocular hypertension model was performed as previously reported (45). Briefly; chronic ocular hypertension was induced using a Nylon 8-0 circumlimbal suture on a tapered needle (8-0 sterile microsuture, AROSurgical Instruments, California, USA) passed subconjunctivally 1.5 mm posterior to the limbus under intraperitoneal Ketamine-Xylazine anesthesia. After making 5-6 sequential subconjunctival passes all around the limbus, the suture was tied off using a slip knot anchored by three simple knots. The suture was left snug, taking care not to directly induce elevated IOP secondary to a tight suture. The sutures then were allowed to slowly tighten over time, resulting in gradual elevation of IOP. In all experiments, both eyes of each animal were subjected to the same treatment to avoid potential confounding contralateral effects.

### Intraocular pressure measurement

A Tonolab rebound tonometer (Icare, Finland) was used to measure the IOP according to the manufacturer’s directions. For each measurement, the tonometer tip was aligned perpendicular to the central cornea. Measurements were obtained at baseline prior to suturing or treatments, following one week of prior alternate day measurements to familiarize the animal to the procedure. Measurements were obtained while the animal was awake between 11 am and 1 pm. Care was taken not to stress the animal or exert pressure on the periocular region during the IOP recordings. Each measurement with the Tonolab rebound tonometer itself consists of six separate readings, of which the highest and lowest are automatically excluded and the mean of the four middle readings are displayed as the final result by the device. For each animal and IOP monitoring session the mean of two consecutive measurements was recorded if they were within 2 mmHg of each other; if there was more than a 2-mmHg difference, then the median of three measurements was recorded.

### Pathological analyses and staining

After euthanasia, eyes were fixed in 4% paraformaldehyde, equilibrated in 30% sucrose, embedded in optimal cutting temperature compound and cryosectioned. 12-μm sections were blocked with 5% donkey serum and probed with primary antibodies to 5-LOX (Novus Biologicals, Catalog # NB110-58748) and 15-LOX (Santa Cruz, Cat. No. sc-133085) according to standard protocols. The sections were washed with PBS-Tween and incubated with fluorescent-conjugated secondary antibodies (Molecular Probes) and DAPI. Subsequently, sections were mounted using MOWIOL 4-88 (Millipore Sigma). Immunofluorescent images were acquired with a Nikon Eclipse-Ti confocal microscope and analyzed with NIS Elements software version 4.51.

### Angle tissue dissection and homogenization

Dissection of a 1-mm strip of rat angle tissue containing a small rim of overlying sclera and cornea, trabecular meshwork, peripheral iris and ciliary body with ciliary processes was carefully performed using Vannas scissors and atraumatic fine forceps. The collected tissue sample was homogenized in aliquoted microfuge tubes, and then snap frozen at −80 °C. Samples were then submitted to quantitative multiplex laser bead analyses (Bio-Plex 200) for assessment of a 27- plex rat cytokine panel and a 3-plex TGF-β panel (Eve Technologies) or lipidomic analysis.

### Statistics

For all experiments, *n* refers to the number of eyes or biological replicates. Graphpad Prism 8.4.3 was used to generate graphs. IOP trend comparisons and lipidomic profile comparisons between two groups were performed using the unpaired t-test. Comparisons between more than two groups were performed using one-way ANOVA with Tukey’s post-hoc analyses. A p-value of less than 0.05 was considered statistically significant.

## RESULTS

### The AA-lipoxin pathway is specifically elevated in glaucomatous aqueous humor

Aqueous humor samples collected from patients with primary open angle glaucoma, or matched controls, were analyzed by targeted LC-MS/MS-based lipidomics. Fifteen patients were enrolled in each group (providing 16 and 18 eye samples in the glaucoma and control groups, respectively). There was no statistically significant difference in age or sex between the two groups (p=0.25 and 0.30, respectively, Table 2). Most patients in the glaucoma group had advanced disease, reflected in a significantly increased cup-to-disc ratio and reduced retinal nerve fiber layer (RNFL) thickness compared to control patient samples (Table 2, Figure 1A, B).

**Figure 1:**
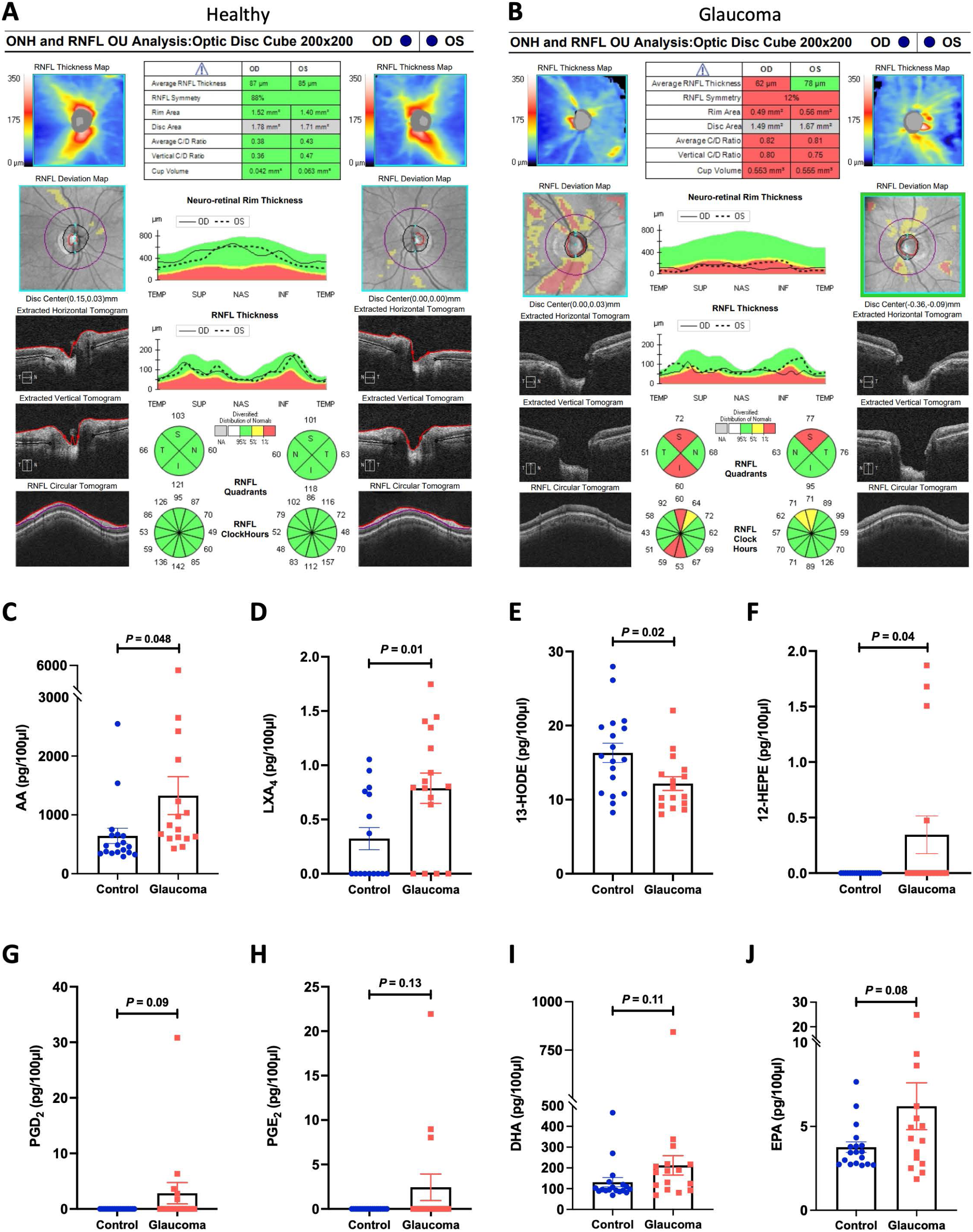
The arachidonic acid-lipoxin pathway is specifically elevated in glaucomatous aqueous humor. **(A)** Representative OCT scans from a control patient showing healthy RNFL and optic nerve head in both eyes. **(B)** Representative OCT scans from a glaucomatous patient showing significant superior and inferior RNFL thinning in the right eye and superior RNFL thinning in the left eye. **(C-F)** Lipidomic analysis of mediators and metabolites from glaucomatous and healthy aqueous humor showed significantly elevated concentrations of **(C)** AA and **(D)** LXA_4_. **(E)** In comparison13-HODE was detected at significantly lower levels in glaucomatous aqueous humor. **(F)** 12-HEPE levels were significantly elevated in the glaucoma group, though statistically driven by only four samples (p values are indicated, bars are SE). **(G-H)** Concentrations of additional analytes detected in human aqueous humor samples included (G) PGD2, (H) PGE2, (I) DHA, and (J) EPA. However, none of these differences reached statistical significance. (For all charts p values are indicated, bars are SE). (OD; right eye, OS; left eye, RNFL; retinal nerve fiber layer).

**Table 2:** Demographics of enrolled patients. There was no significant difference between the two groups for age and sex. IOP was similar between the two groups as advanced glaucomatous eyes were treated medically to achieve a low target pressure. The cup-to-disc ratio was higher for glaucomatous eyes, reflective of the advanced disease stage.

|  | **Glaucoma** | **Control** | **p value** |
| --- | --- | --- | --- |
| Number of eyes (patients) | 16 (15) | 18 (15) | -- |
| Age | 68.7 ± 6.4 years | 71.0 ± 4.7 years | 0.25 |
| Sex | 9 males, 7 females | 6 males, 12 females | 0.3 |
| Intraocular pressure | 14.1 ± 3.1 mmHg | 15.2 ± 1.6 mmHg | 0.24 |
| Cup-to-disc ratio | 0.9 ± 0.1 | 0.3 ± 0.1 | <0.001 |

Lipidomic analysis focused on lipoxygenase (LOX) and cyclooxygenase (COX) pathways and their polyunsaturated fatty acid (PUFA) substrates. The analysis included eicosanoids (prostaglandins, leukotrienes, lipoxins), AA and other PUFA, pathway markers and metabolites, and select ω-3 PUFA derived SPMs (see Supplementary Table 1 for a full list of these results).

Four mediators showed striking changes in concentration between groups. Most significant were the substrate AA (643.07 ± 127.15 vs 1328.04 ± 312.43 pg/100 μl of aqueous humor, p=0.048) and the AA lipoxygenase product LXA_4_ (0.74 ± 0.08 vs 1.05 ± 0.36, p=0.01), whose concentrations were strongly elevated in the aqueous humor of glaucoma patients (Figure 1C, D). In addition, 13-HODE (13-hydroxyoctadecadienoic acid), a product of linoleic acid lipoxygenase metabolism by 15-LOX, was present at significantly lower levels in glaucomatous aqueous humor (Figure 1E). The levels of 12-HEPE (12-hydroxyeicosapentaenoic acid), an EPA-derived metabolite, were also significantly higher in glaucomatous aqueous humor (1.38 ± 0.62 vs 0 pg/100 μL, p=0.04), but this result was less convincingly driven by changes in only a few samples (Figure 1F). In addition, the AA-derived prostaglandins PGE_2_ (Prostaglandin E_2_) and PGD_2_ (Prostaglandin D_2_) were elevated in some glaucoma samples, as well as the ω-3 PUFAs docosahexaenoic acid (DHA), and eicosapentaenoic acid (EPA), but these changes did not reach significance between the groups (Figure 1G-H).

No other PUFA, LOX or COX metabolites were identified at significant levels by our LC-MS/MS method (Supplementary Table 1). Therefore, these results suggest a select response in the activity of AA and LXA_4_ circuits that generates a marked increase in their levels in glaucomatous aqueous humour. Formation and increased levels of LXA_4_ are consistent with immunofluorescent analysis of anterior segment outflow tissues from healthy human donor eyes and glaucomatous patient eyes. Ciliary muscle, vasculature and outflow tissues demonstrated consistent staining for the LXA_4_ biosynthetic enzymes 5-LOX and 15-LOX (Supplementary Figure 2). Interestingly, matched sections from two glaucomatous patient eyes showed prominently increased 5-LOX staining and mildly increased 15-LOX staining in the trabecular meshwork (Supplementary Figure 2).

### The AA-lipoxin circuit is induced by latanoprost and timolol treatment in human TM cells

We had predicted that lipoxin levels would be reduced in glaucoma patients due to their well-documented pro-resolution activities, and from reports of other chronic diseases which exhibit this pattern (46–50). Therefore, the marked elevation of LXA_4_ we observed in glaucomatous aqueous fluid was unexpected. Upon consideration, we wondered whether this response might be caused by the topical IOP-lowering medications the patients were taking. In particular, the most commonly prescribed glaucoma drug, latanoprost (6–8), is a prostaglandin F_2_α (PGF_2_α) analogue that could potentially impact LOX or COX pathways and production of other AA metabolites. Notably, all enrolled glaucoma patients were taking topical glaucoma medications, and usually more than one; with 100% taking PGF_2_α analogues, 93.75% on beta blockers, 87.5% on carbonic anhydrase inhibitors, and 75% taking alpha-2 adrenergic agonists (Figure 2A). Therefore, a majority of patients were typically prescribed combinations of these drug classes.

**Figure 2:**
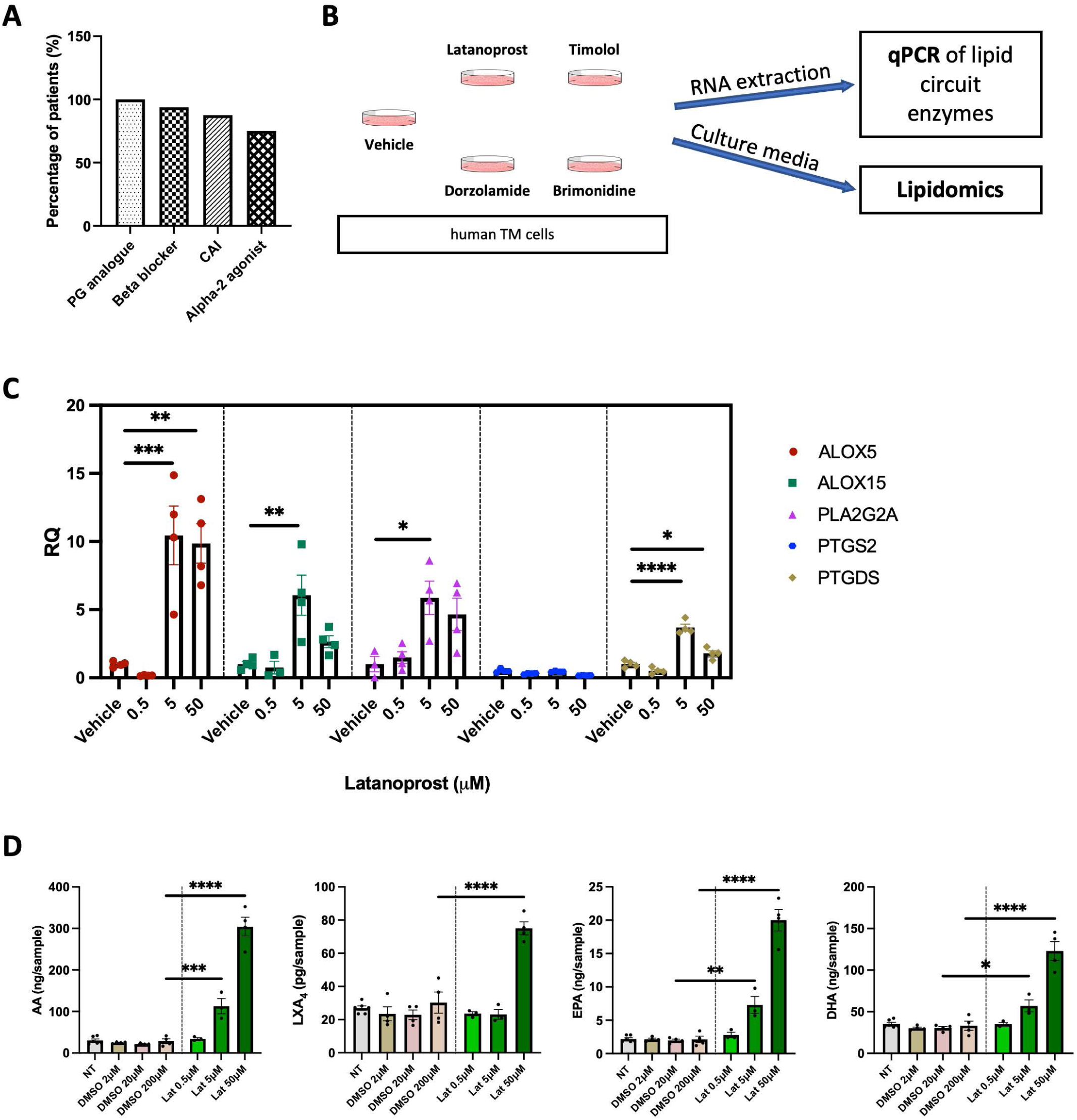
The AA-lipoxin circuit is induced by latanoprost and timolol treatment in human trabecular meshwork cells. **(A)** Graph representing the percentage of glaucoma patients taking topical glaucoma eye drops, including prostaglandin analogues, beta blockers, carbonic anhydrase inhibitors and alpha-2 adrenergic agonists. **(B)** Human trabecular meshwork cells were treated with latanoprost (prostaglandin analogue), timolol (beta blocker), dorzolamide (carbonic anhydrase inhibitor), brimonidine (alpha-2 adrenergic agonist), or vehicle for one hour before collecting RNA for qPCR and conditioned media for lipidomic analyses. **(C)** Quantification of qPCR results showed that treatment with latanoprost caused a significant, dose-dependent upregulation of ALOX5, ALOX15, PLA2G2A and PTGDS expression. **(D)** Lipidomic analyses of the culture media showed a significant increase in arachidonic acid and LXA_4_ levels with increasing latanoprost treatment. In addition, EPA and DHA substrate levels were also significantly elevated by treatment compared to vehicle. (****p<0.0001, ***p<0.001, **p<0.01, *p<0.05, ns; not significant, bars are SE) (AA; arachidonic acid, ALOX5; arachidonate 5-lipoxygenase, ALOX15; arachidonate 15-lipoxygenase, DHA; docosahexaenoic acid, EPA; eicosapentaenoic acid, LXA_4_; lipoxin A_4_, PLA2G2A; phospholipase A_2_ group IIA, PTGDS; prostaglandin D_2_ synthase, PTGS2; prostaglandin-endoperoxide synthase 2 (cyclooxygenase-2), RQ; relative quantification).

In order to test the potential influence of these medications on the AA-lipoxin circuit, an *in vitro* experiment was designed in which human trabecular meshwork (TM) cells were directly treated with clinically relevant concentrations of common drugs representing each of the four classes, or vehicle, and analyzed for changes in eicosanoid and lipoxin pathways. After one hour, RNA was isolated from the treated cells to assess expression of relevant enzymes using quantitative reverse-transcription polymerase chain reaction (qPCR). In parallel, conditioned media was also collected for corresponding lipidomic analyses (Figure 2B).

Interestingly, latanoprost treatment significantly induced dose-dependent expression of key enzymes for generating lipoxins. Expression of the rate-limiting enzyme, 5-LOX (ALOX5), was markedly upregulated by 10.45 and 9.86-fold (p=0.001) in human TM when treated with 5 μM and 50 μM, respectively, compared to vehicle treatment (Figure 2C). Similarly, the second required enzyme for lipoxin formation 15-LOX (ALOX15), was upregulated 6.06-fold by 5 μM latanoprost (p=0.003). The enzyme that generates AA from phospholipids, phospholipase A_2_ (PLA2G2A) was also upregulated 14.23-fold by latanoprost at 5 μM (p=0.015). Latanoprost did not induce expression of COX-2 (PTGS2), but increased expression of prostaglandin D_2_ synthase (PTGDS), which was upregulated 3.68 (p<0.0001) and 1.79 (p=0.02) fold by 5 μM and 50 μM latanoprost treatment, respectively (Figure 2C).

In parallel, conditioned media samples from this experiment were also analyzed by lipidomics for functional changes in lipoxin pathways. Consistent with the increases observed in PLA2G2A, 5-LOX and 15-LOX gene expression, corresponding AA and LXA_4_ levels were significantly elevated in latanoprost treated samples in a dose-dependent manner (Figure 2D). Compared to vehicle, 5 and 50 μM latanoprost treatment caused 5.32 and 10.7-fold increased levels of AA (p=0.0002 and p<0.0001, respectively), and LXA_4_ levels were significantly increased (2.48 fold) with 50 μM latanoprost (p<0.0001). Interestingly, levels of additional substrates that are released by PLA_2_ were also increased, including EPA (5 and 50 μM latanoprost; p=0.002 and p<0.0001, respectively), and DHA (5 and 50 μM latanoprost; p=0.002 and p<0.0001, respectively; Figure 2D).

Of the other drug classes tested, a somewhat mixed picture is presented. Surprisingly similar trends to latanoprost were observed in cells treated with the beta-adrenergic receptor antagonist timolol; ALOX5 was upregulated 8.12-fold (p<0.0001) by 100 μM timolol treatment compared to vehicle. ALOX15 was upregulated 3.22, 3.23 and 5.98-fold by 1, 10 and 100 μM timolol, respectively (p=0.03, 0.03 and <0.0001; Supplementary Figure 3A). PLA2G2A was upregulated 16.34, 11.82 and 11.79-fold, respectively by 1, 10 and 100 μM timolol, respectively (p=0.0002, 0.003 and 0.003, respectively), and PTGDS was upregulated 4.49, 5.29 and 10.28-fold by 1, 10 and 100 μM timolol, respectively (p=0.007, 0.002 and <0.0001, respectively; Supplementary Figure 3A). In contrast, neither treatment with the carbonic anhydrase inhibitor dorzolamide, or alpha-2 agonist brimonidine, showed substantial effects on expression of the same gene panel (Supplementary Figure 4A, B). Consistent with the qPCR results, treatment with 10 and 100 μM timolol significantly elevated the levels of AA (16.46 and 19.37 fold, respectively; p<0.0001 for both), and 100 μM timolol elevated levels of LXA_4_ (1.74 fold, p=0.03; Supplementary Figure 3B). Concentrations were also elevated at 10 and 100 μM for EPA (p<0.0001) and DHA (p<0.0001; Supplementary Figure 3B). Dorzolamide or brimonidine did not generate substantial or consistent changes in AA, EPA or DHA. Although, LXA_4_ levels were mildly increased with 0.5 and 5 μM treatment (p=0.0004 and 0.04, respectively; Supplementary Figure 4C). Treatment with 200 μM brimonidine significantly increased levels of AA, EPA and DHA (p<0.0001 for all) but did not result in a significant increase in LXA_4_ levels (Supplementary Figure 4D).

Together, these results present a picture that is fairly consistent with the patient lipidomic data, indicating that latanoprost treatment, and to a lesser extent timolol, specifically amplifies the AA-lipoxin synthetic circuit in TM cells.

### The AA-lipoxin circuit is not induced by ocular hypertension alone

Based on our clinical results, an alternative possibility is that ocular hypertension alone induces the AA-LXA_4_ pathway. Yet, it was not possible to acquire untreated clinical aqueous humour samples from glaucoma patients due to ethical considerations. Therefore, in order to test this possibility we turned to a recently characterized rat model of gradual ocular hypertension (gOHT) generated by slack circumlimbal sutures that tighten over time (45). Once elevated, ocular hypertension was maintained for 8 weeks in the sutured eyes, at which time they were processed for lipidomic analyses of angle tissues (Figure 3C, D). In these OHT samples AA concentrations were not significantly different from control normotensive eyes, and the trend was towards reduced levels (Figure 3E). Likewise, levels of prostaglandins PGE_2_, PGD_2_, and 6-keto-PGF_1α_ were detected, but were not significantly altered and exhibited a similarly reduced trend (Figures 3F-H). Notably, LXA_4_ levels analyzed from each single eye were below the detection threshold for all groups, which is likely due to the small tissue sample sizes. These findings contrast with the clinical lipidomic results and indicate that ocular hypertension by itself does not increase AA metabolism *in vivo*.

**Figure 3:**
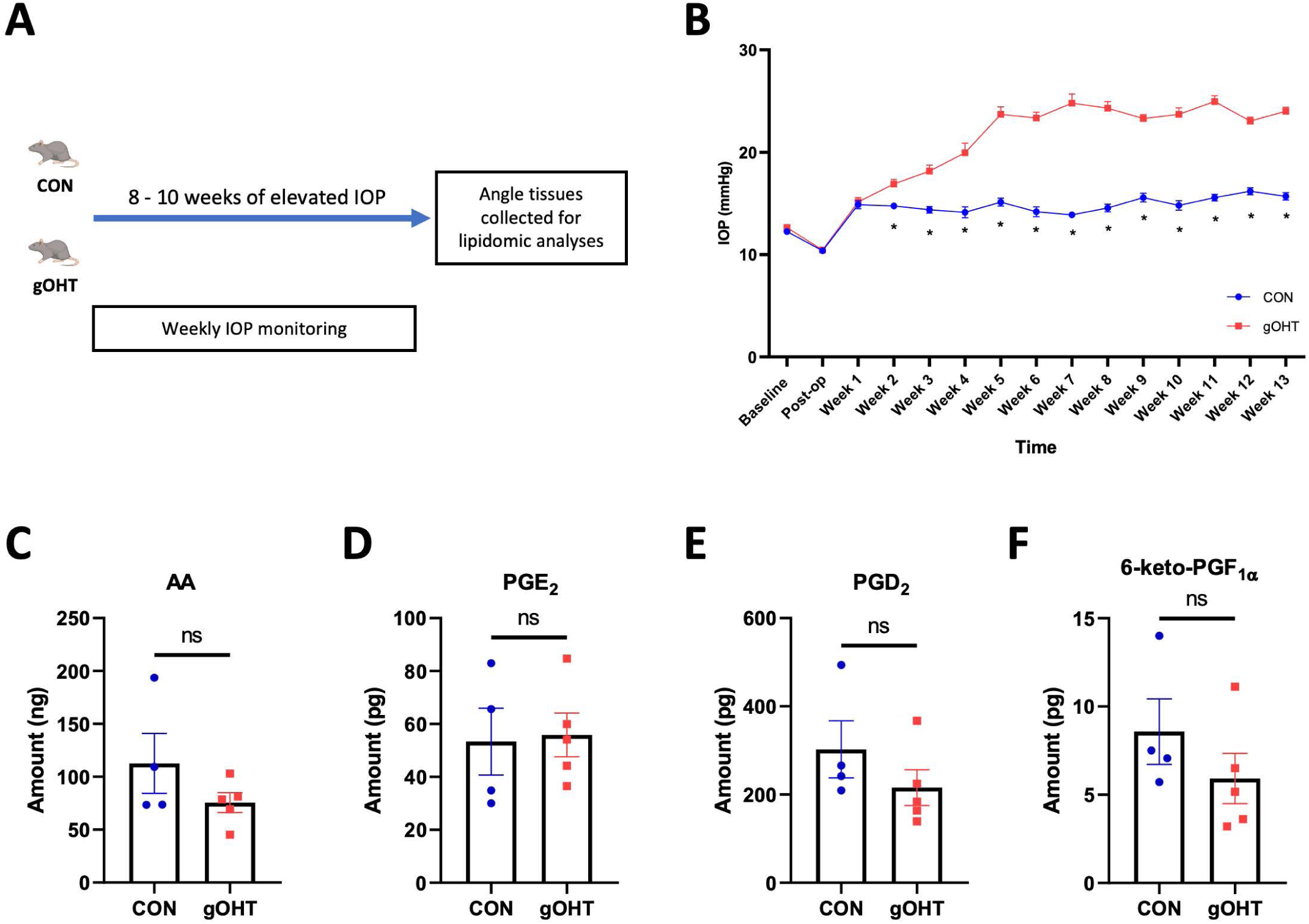
The AA-lipoxin circuit is not induced by ocular hypertension alone. **(A)** Gradual ocular hypertension (gOHT) was induced by a circumlimbal suture in six-week-old Long Evans rats and maintained for 8-10 weeks before eyes were collected for lipidomic analyses. **(B)** As expected, circumlimbal suturing induced a gradual increase in IOP, consistently exceeding 20 mmHg from weeks 3-5 post-suturing (*p<0.0001, bars are SE). **(C)** Lipidomic analyses detected AA levels that were not significantly altered in the OHT group compared to control. **(D-F)** Similarly, endogenous prostaglandins D_2_ and E_2_, and 6-keto-prostaglandin F_1α_ were detected, but were not significantly altered by ocular hypertension alone (bars are SE). (PGD_2_, prostaglandin D_2_; PGE_2_, prostaglandin E_2_).

In contrast, topical administration of latanoprost or vehicle to rat eyes resulted in lipid mediator profiles that largely overlapped with the clinical and human cell culture results. Rats were dosed daily with 40 μL of latanoprost (0.005%) for seven days, and the angle tissues collected for lipidomic analyses (Figure 4A). In this case the sample homogenates (n=8) were pooled to increase levels and improve detection of intermediates and LOX products in these small rat tissue samples. The AA and DHA PUFA substrates were slightly reduced by latanoprost treatment (Figure 4B). More importantly, similar to the clinical and human TM cell samples, levels of LXA_4_ were elevated, along with several key pathway intermediates and products of the lipoxin biosynthetic pathway, in latanoprost treated samples compared to vehicle controls (Figure 4C). Interestingly, a panel of intermediates and products of the cyclooxygenase (COX) pathway were also elevated, including PGE_2_ and PGD_2_, as in the clinical samples (Figure 4D). Finally, LTB_4_, a 5-LOX product, was sharply reduced (Figure 4E) in sharp contrast to the increase in LXA_4_; a pattern consistent with a shift of 5-LOX activity to generating proresolving mediators instead of pro-inflammatory mediators (51, 52).

**Figure 4:**
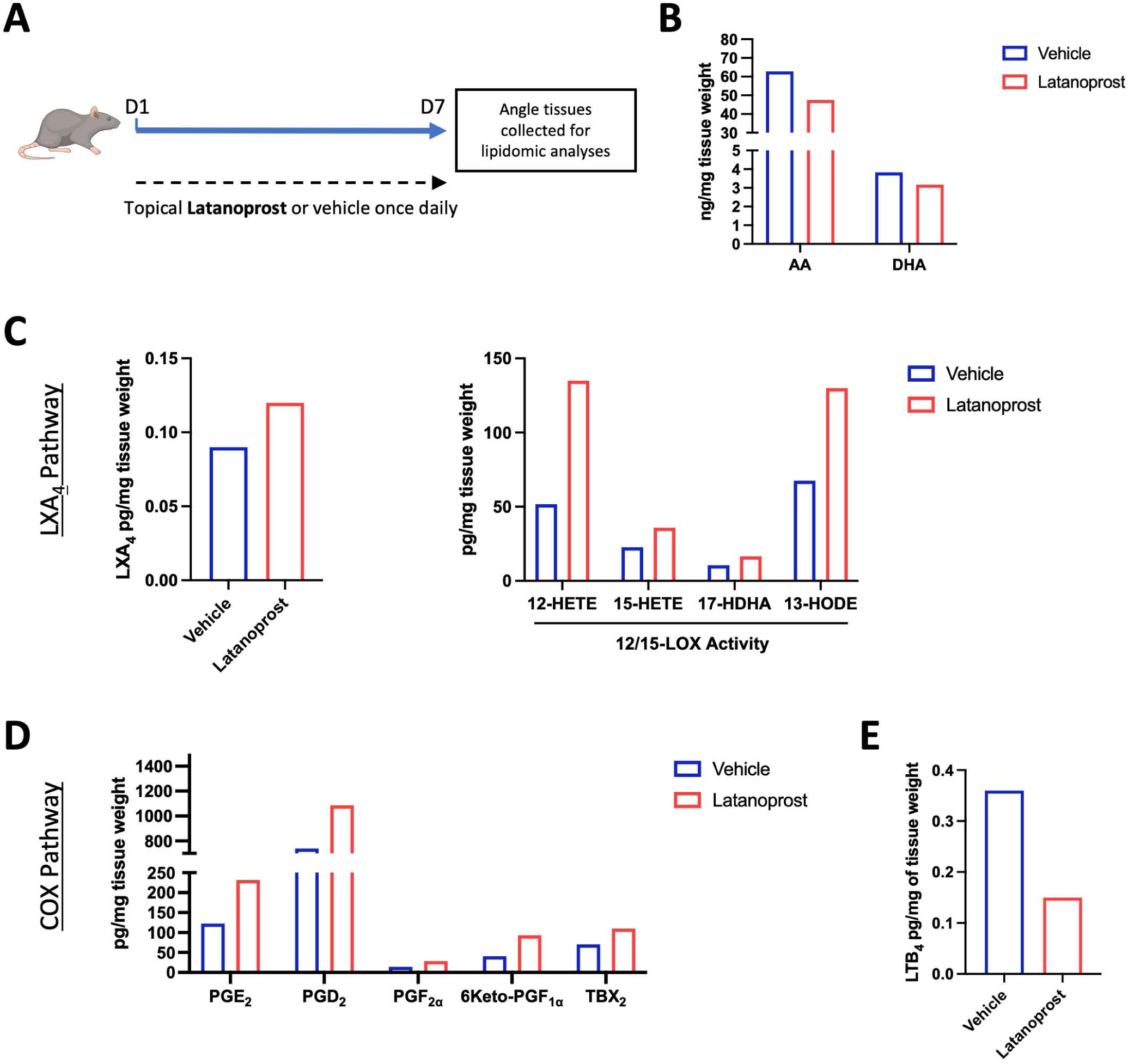
The LXA_4_ pathway and cox pathways are induced by latanoprost treatment *in vivo*. **(A)** Six-week-old Long Evans rats were administered topical latanoprost for 7 days, followed by the analyses of angle tissues. **(B)** Concentrations of AA and DHA were slightly reduced in latanoprost treated samples compared to vehicle (bars are composites of 8 samples). **(C)** Elevated levels of LXA_4_ were detected, along with several pathway intermediates and products, including 12-HETE, 15-HETE, 17-HDHA, and 13-HODE (bars are composites of 8 samples). (**C**) Elevated cox pathway products were also detected, including PGE_2_, PGD_2_, PGF_2α_, 6Keto-PGF1α, and TBX2 (bars are composites of 8 samples). (**D**) Levels of LTB4 were sharply reduced in latanoprost treated samples compared to vehicle (bars are composites of 8 samples).

In comparison, Timolol was also administered topically with the same experimental design, and generally resulted in no substantial changes in LXA_4_ or COX pathway products in the rat model (Supplementary Figure 4). Together these data provide direct evidence that latanoprost specifically promotes AA metabolism and LXA_4_ synthesis, as well as prostaglandin production *in vivo*.

### LXA_4_ does not cause acute IOP-lowering but inhibits proinflammatory cytokines and induces production of TGF-β3

To study the effect of elevated LXA_4_ itself on IOP and the outflow tissues, six-week-old Long Evans rats were treated topically by eye drop with 40 μM LXA_4_, once daily, for seven days (Figure 5A). IOP was monitored during the first 24 hours and then daily till the end of the experiment. Over this period there was no significant difference in IOPs between LXA_4_ treated and vehicle treated eyes (Figure 5B). This indicates that LXA_4_ alone is not sufficient to reduce IOP.

**Figure 5:**
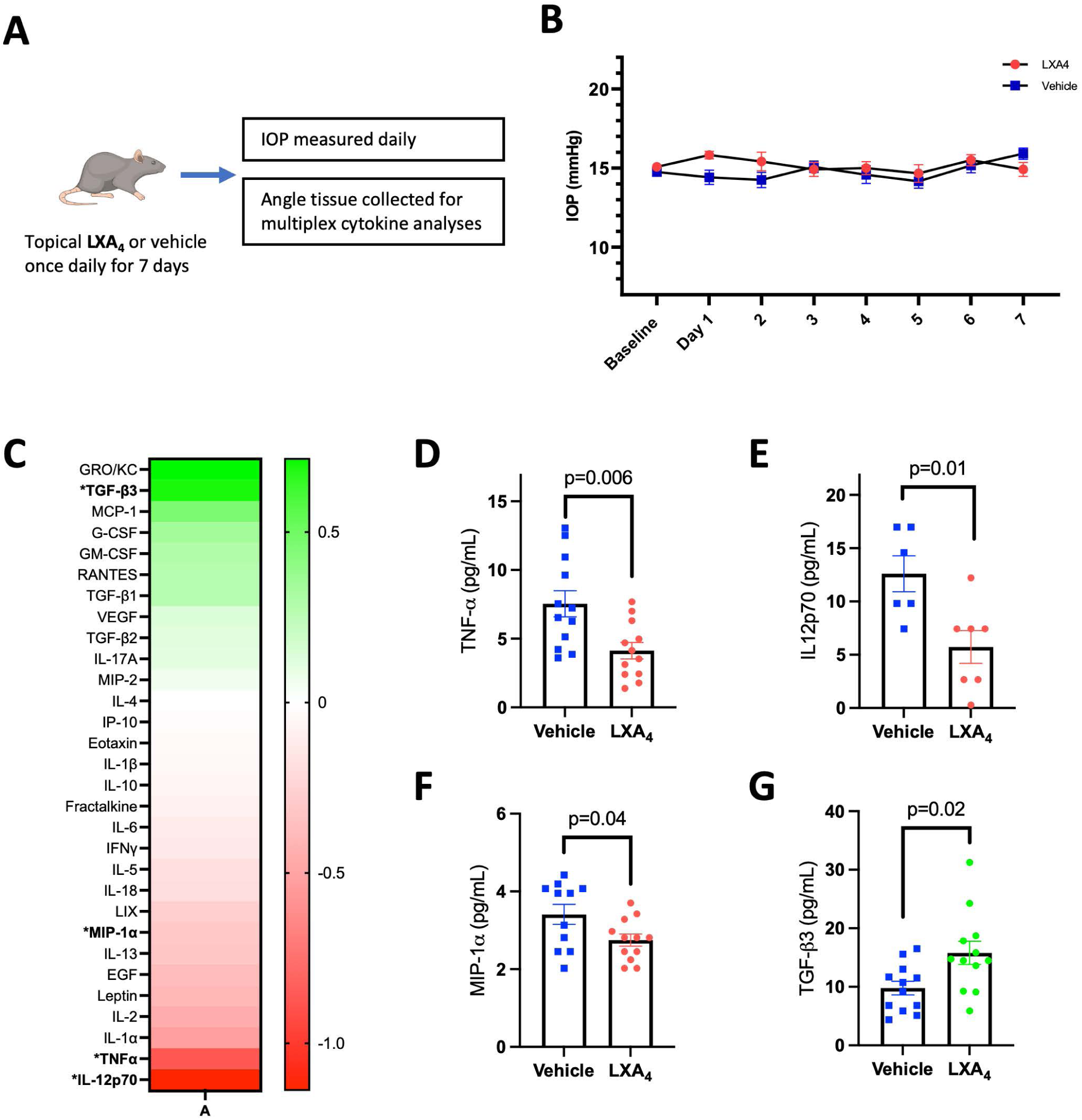
LXA_4_ does not cause acute IOP-lowering, but inhibits proinflammatory cytokines and induces production of TGF-β3. **(A)** Six-week-old Long Evans rats were treated with LXA_4_ once daily for one week, and the eyes were collected for cytokine analyses of the angle tissues. **(B)** Daily IOP measurements indicate that LXA_4_ did not cause significant IOP changes compared to vehicle treated controls (bars are SE). **(C-F)** At the end of the study angle tissue samples were subjected to a panel of 30 cytokines, with those showing significant difference presented here. There was a significant decrease in the levels of pro-inflammatory cytokines **(C)** IL-12, **(D)** MIP-1α and **(E)** TNF-α, and **(F)** a significant increase in TGF-β_3_ levels (p values indicated, bars are SE). (IL-12p70; interleukin-12p70, MIP-1α; macrophage inflammatory protein-1 alpha, TGF-β_3_, transforming growth factor-beta_3_, TNF-α; tumor necrosis factor-alpha).

However, as LXA_4_ has potent anti-inflammatory and proresolution activities (46, 53, 54), we also profiled whether repeated treatment would alter inflammation signaling in outflow tissues. Angle tissues were harvested from vehicle and LXA_4_-treated rat eyes and subjected to a cytokine panel of 30 mediators (Supplementary Table 2). Significantly altered cytokines were interleukin-12 (IL-12; 12.60 vs 5.73 pg/mL, p=0.01), macrophage inflammatory protein-1α (MIP1α; 3.41 vs 2.75 pg/mL, p=0.04) and tumor necrosis factor-alpha (TNF-α; 7.55 vs 4.12 pg/mL, p=0.006), whose levels were all significantly lower in LXA_4_-treated eyes (Figure 5C-E). These results are consistent with expected anti-inflammatory effects. In comparison, transforming growth factor-β_3_ (TGF-β_3_) concentrations were significantly higher in LXA_4_-treated eyes compared to vehicle-treated eyes (15.79 vs 9.78 pg/mL, p=0.02; Figure 5F). TGF-β_3_is part of the TGF-β superfamily of cytokines that promote extracellular matrix deposition and remodeling, and has been notably linked to TM cells and glaucoma through an extensive literature (55–59).

### Prostaglandin synthesis is required for latanoprost IOP-lowering activity

As LXA_4_ did not mediate acute IOP-lowering, we wondered whether this component of latanoprost activity might be generated by another branch of AA metabolism to generate an autocrine cycle of prostaglandin synthesis. Therefore, we sought to block prostaglandin production in the context of latanoprost treatment. The two COX enzymes, COX-1 and COX-2, catalyze the formation of prostaglandins from AA (60). Bromfenac preferentially inhibits COX-2, although it also targets COX-1, and demonstrates potent anti-inflammatory effects by blocking prostaglandin synthesis (61). Bromfenac is widely used in the eye after cataract surgery to decrease the risk of cystoid macular edema secondary to ocular inflammation (62). Rat eyes were treated with topical bromfenac daily for two days before initiating daily latanoprost eye drops for one week. In select groups bromfenac administration was continued during the seven days of latanoprost treatment (Figure 6A). Lipidomic analyses of angle tissues showed strong inhibition of prostaglandin synthesis by bromfenac (Figure 6B). Upon IOP measurement, eyes treated with bromfenac alone showed no IOP change compared to vehicle. As expected, treatment with latanoprost alone significantly lowered IOP. However, eyes treated with both bromfenac and latanoprost exhibited a significantly reduced IOP decrease compared to latanoprost alone (Figure 6C). Quantification of average IOP between days 2-7 (when latanoprost showed maximal IOP lowering), revealed a significant difference between latanoprost treatment alone, and cotreatment with bromfenac and latanoprost (p<0.001, Figure 6D). These findings indicate that synthesis of endogenous prostaglandins is required for full IOP-lowering actions of latanoprost.

**Figure 6:**
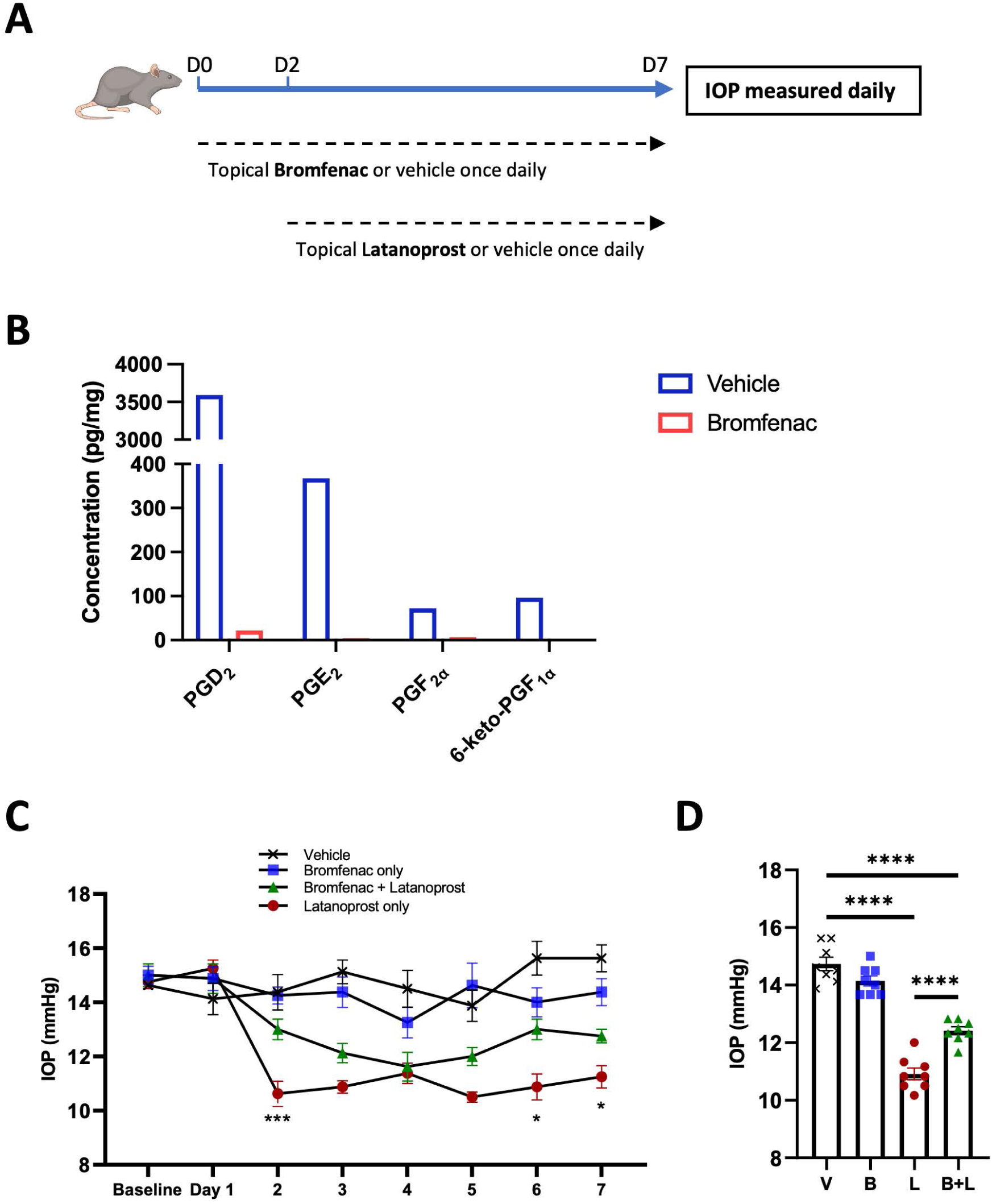
Prostaglandin synthesis is required for latanoprost acute IOP-lowering activity. **(A)** Six-week-old Long Evans rats were treated with the COX inhibitor, bromfenac, or vehicle for 48 hours prior to starting concomitant latanoprost treatment over the next seven days. Both treatments were administered once daily, with daily IOP monitoring. **(B**) Analyses of rat angle tissues following administration of bromfenac (B) or vehicle (V) showed strong inhibition of levels of the prostaglandins PGD_2_, PGE_2_, PGF_2α_, and 6-keto-PGF_1α_ (bars are composites of 6 samples). (**C**) Bromfenac treatment alone had no effect on IOP, while latanoprost treatment caused a rapid and sustained IOP reduction. When administered together, bromfenac treatment attenuated the IOP-lowering activity of latanoprost (***p<0.001, *p<0.05 between latanoprost and latanoprost + bromfenac, bars are SE). **(D)** Comparison of the average IOP from days 2 to 7 demonstrated significant attenuation of the IOP-lowering effect of latanoprost by bromfenac (****p<0.0001, bars are SE). (COX; cyclooxygenase).

## DISCUSSION

The production and roles of lipid mediators derived from PUFA through the LOX and COX pathways remain surprisingly unexplored in glaucoma patients. In our study, aqueous humor lipidomic analyses of COX and LOX derived mediators showed a strong and selective upregulation of the AA-LXA_4_ pathway in glaucoma patients compared to non-glaucomatous controls. This was a strikingly selective upregulation, considering the panel of 40 mediators and intermediates assessed, including the ω-3 substrates DHA and EPA, along with a variety of active metabolites. This robust increase in LXA_4_ production in patients was surprising, given its established protective and anti-inflammatory actions. Yet, LXA_4_ formation can be promoted pharmacologically in other contexts (33–38). Since all glaucoma patients included in this study were unavoidably taking topical IOP lowering medications, including prostaglandin mimetics, we evaluated their direct effect on the AA-LXA_4_ pathway. Together, our results indicate that latanoprost induces a dose-dependent increase in LXA_4_ production *in vitro*, and *in vivo*. This result is supported by an early study in human blood leukocytes that had reported the ability of another prostaglandin, PGE_2_, to induce lipoxin formation *de novo* by regulating 15-LOX expression (51). In contrast, elevated IOP alone did not activate this pathway *in vivo*. Exogenous LXA_4_ in turn had no acute effect on IOP, but strongly inhibited proinflammatory cytokines and stimulated production of TGF-β_3_. In a parallel pathway, COX-mediated prostaglandin synthesis was required for the acute IOP-lowering effects of latanoprost. Together, these results suggest a new model for parallel acute and long-term latanoprost mechanisms that can be uncoupled to involve either prostaglandin or lipoxin actions, respectively (Figure 7).

**Figure 7:**
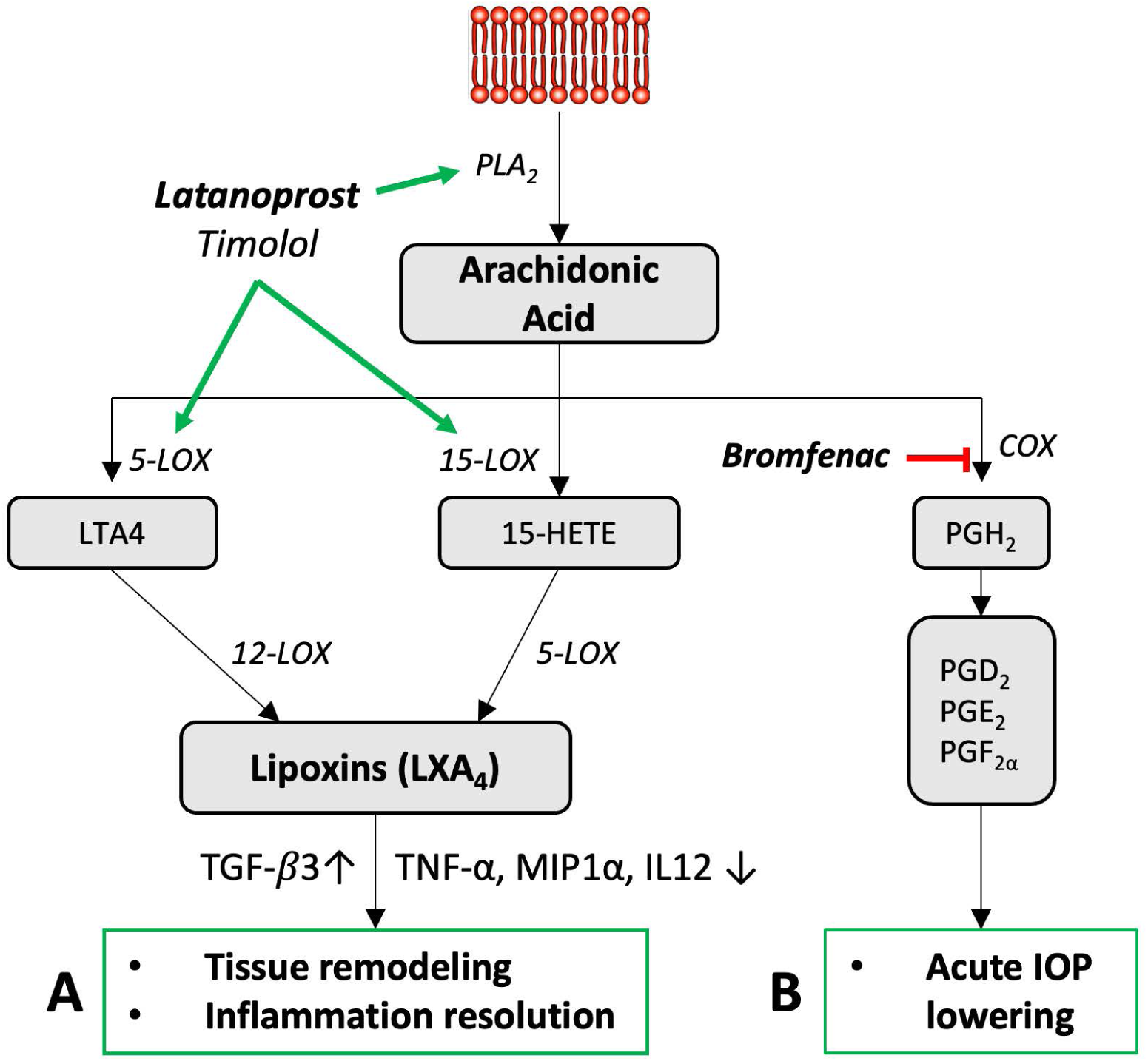
Flowchart depicting a proposed parallel AA-dependent drug mechanism. **(A)** The upregulation of PLA_2,_ 5-LOX and 15-LOX by latanoprost induces synthesis of LXA_4_, resulting in tissue remodeling and inflammation resolution to exert sustained IOP lowering effects. (**B**) In concert, endogenous prostaglandins are generated via COX activity, resulting in acute IOP lowering. (PGD_2_; prostaglandin D_2_, PGE_2_; prostaglandin E_2_, PGF_2α_; prostaglandin F_2α_).

Prostaglandin analogues are widely used as the first-line treatment for open-angle glaucoma due to their once-daily dosing regimen and substantial IOP reduction (6–8, 63). In fact, latanoprost alone is one of the most commonly prescribed medications, with nearly 10 million prescriptions in the U.S. in 2021 (https://clincalc.com/DrugStats/Drugs/Latanoprost). Yet, as a class the mechanisms underlying these drug actions are still unclear. Latanoprost is an analogue of PGF_2α_, with an isopropyl ester substituent replacing the α-carboxylic acid. It is thought to lower IOP by increasing the outflow of aqueous humor through the uveoscleral (10) and trabecular meshwork pathways (64, 65). The established mechanism of action of latanoprost involves binding to a G-protein coupled FP receptor, which is expressed in the ciliary muscle and the trabecular meshwork of the eye (12). Traditionally, activation of the FP receptor by prostaglandin F_2α_ analogues is thought to stimulate phospholipase A2, resulting in the release of arachidonic acid (AA) and subsequent synthesis of endogenous prostaglandins, including PGE_2_. Production of PGE_2_ induces cAMP that promotes smooth muscle relaxation to enhance aqueous humor outflow and reduce IOP. Short-term treatment in primates with PGF_2α_ results in rapid IOP reduction, and normalization after cessation of treatment (66–68). We observed some increased prostaglandin synthesis following latanoprost treatment. Also, inhibition of endogenous prostaglandin production by blocking COX activity significantly reduced the acute IOP actions of latanoprost. These results are consistent with a pseudo-autocrine loop contributing to acute IOP lowering, as previously proposed (69). Accordingly, caution has been suggested when prescribing topical NSAIDs to glaucoma patients using prostaglandin analogues (70), although conflicting results have also been reported (71), and our results suggest more research in this area is needed.

In addition to mediating acute, but transient, IOP-lowering activities, increasing evidence describes how prostaglandin analogues also remodel the extracellular matrix of the trabecular meshwork and ciliary body. This activity occurs via secretion of MMPs-1, 2, 3 and 9, increasing the permeability of these tissues to aqueous humor (16, 72). Interestingly, long-term treatment with these agents results in long-lasting IOP reduction that persists even after treatment is stopped (17, 73), which has been partially attributed remodeling in the ciliary body (13, 15). In some patients, treatment with prostaglandin analogues resulted in lowered IOP which persisted even months after cessation of treatment (17). Recently, Park *et al*. reported decreased anterior scleral thickness following prostaglandin analogue treatment, which was linked to a similar mechanism (74). Yet, the mechanism of action that directs this tissue remodeling pathway is not well understood (15). Our results indicate that treatment with latanoprost upregulates 5- and 15-LOX, resulting in increased synthesis of LXA_4_. Supplementation of LXA_4_ resulted in a marked anti-inflammatory effect and increased production of TGF-β_3_, which has been directly linked to trabecular meshwork remodeling (55, 75, 76). Thus, this work has identified a novel branch of the latanoprost signaling mechanism that explains its biochemical connection to ECM remodeling and provides a new link to inflammation resolution (Figure 7).

Interestingly, similar to latanoprost, Timolol also induced upregulation of PLA_2_, 5-LOX and 15-LOX *in vitro*, with increased synthesis of AA and LXA_4_. Beta adrenergic antagonists act primarily through decreased cytosolic cAMP levels and altered calcium signaling (77). Activation of the AA-LXA_4_ pathway requires calcium signaling, which may partially explain the unexpected actions of timolol. However, these timolol results were not repeated in rat eyes *in vivo*. Timolol canonically reduces IOP by decreasing aqueous humor secretion (78), but recently has also been reported to reduce aqueous outflow facility in healthy human eyes through an unknown mechanism (79). Upregulation of AA and its downstream mediators by timolol suggests a potential common or interacting mechanism of action with latanoprost. Yet, the detailed interactions that mediate these effects will require further clarification.

In summary, although their use is widespread, the molecular mechanisms underlying the actions of prostaglandin analogues in the eye have remained unclear. We report an upregulation of AA-LXA_4_ induced by latanoprost that may explain long-term effects. Given the well-established pro-inflammatory roles of PGF_2_α (80), this unanticipated pathway results in pro-resolving, anti-inflammatory and remodeling changes in the outflow tissues that can be uncoupled from its acute IOP-lowering effects. The roles of AA and its LOX products have not been explored before in the context of glaucoma and IOP lowering. Therefore, these insights may provide a foundation for investigating new ocular hypotensive therapeutic targets with sustained anti-inflammatory and remodeling actions. Finally, our findings also suggest a note of caution to carefully interpret similar patient biomarker studies to distinguish observations due to the disease process itself from changes resulting from treatments.

## ACKNOWLEDGEMENTS

Grateful thanks to D. Peters for the gift of TM-1 cells, and B.A. Flitter for help troubleshooting. J. Sivak holds the UHN Foundation Trope Research Chair. D. Mathew held a University of Toronto Vision Science Research Program (VSRP) fellowship. Funding was provided by CIHR grants PJT166201 and PJT168845 (JS), and NIH grant R01EY030218 (JS, JF, KG).

