## Supplementary Figures for "Lipidomic Analysis Reveals Drug-Induced Lipoxins in Glaucoma Treatment"

Supplementary Figure 1

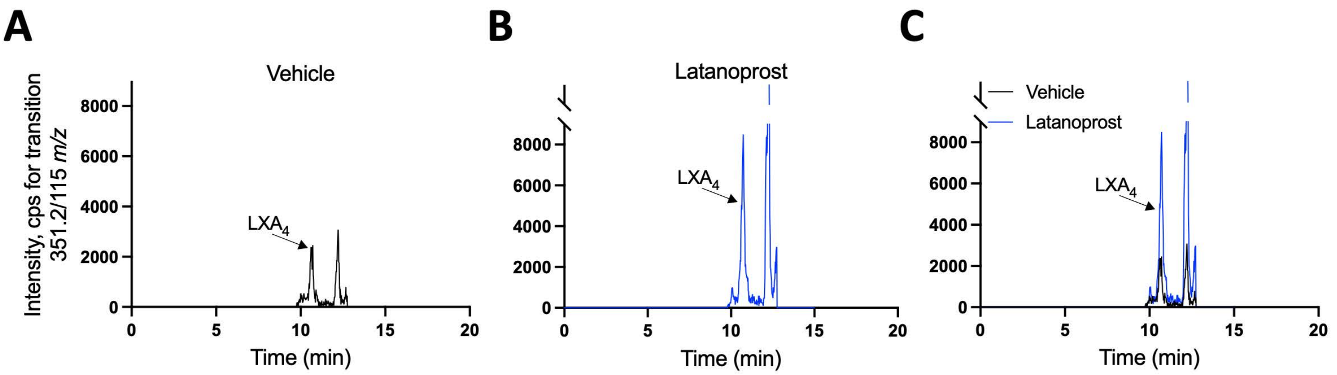


**Supplementary Figure 1. Representative LC-MS/MS analyses in negative ion mode using scheduled Multiple Reaction Monitoring (MRM).** Extracted ion chromatogram (XIC) showing the intensity, counts per second (cps) for $LXA_4$ transition 351.2/115 *m/z* on the Y-axis and retention time on the X-axis for **A)** Vehicle (black), **B)** Latanoprost 50μm (blue), **C)** Combined graph for vehicle (black) and latanoprost 50μm (blue). The arrow indicates the $LXA_4$ peak, which matches the retention time for $LXA_4$ in external standards.

.

Supplementary Figure 2

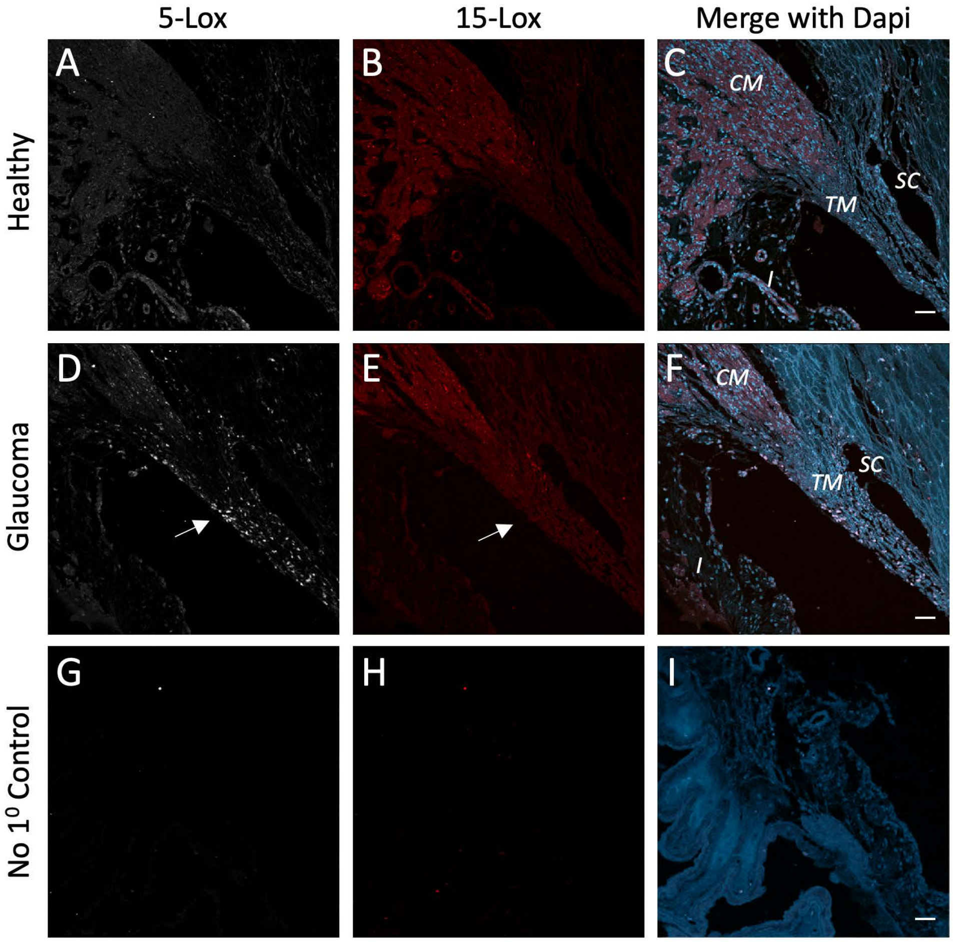


**Supplementary Figure 2. 5- and 15-LOX staining of human anterior segment tissues. (A-C)** Representative sections from a healthy human eye, stained with antibodies directed to 5-LOX (white) and 15-LOX (red), highlight the ciliary muscle (*CM*), iris (*I*) and associated vasculature, and outflow tissues, including the trabecular meshwork (*TM*) and Schlemm's canal (*SC*). **(D-F)** Representative sections from a human glaucomatous eye showing increased 5-LOX, and slightly increased 15-LOX in the *TM* (arrows). **(G-I)** Control sections without primary antibody staining were generally blank. (Scale bars represent 50 μm).

Supplementary Figure 3

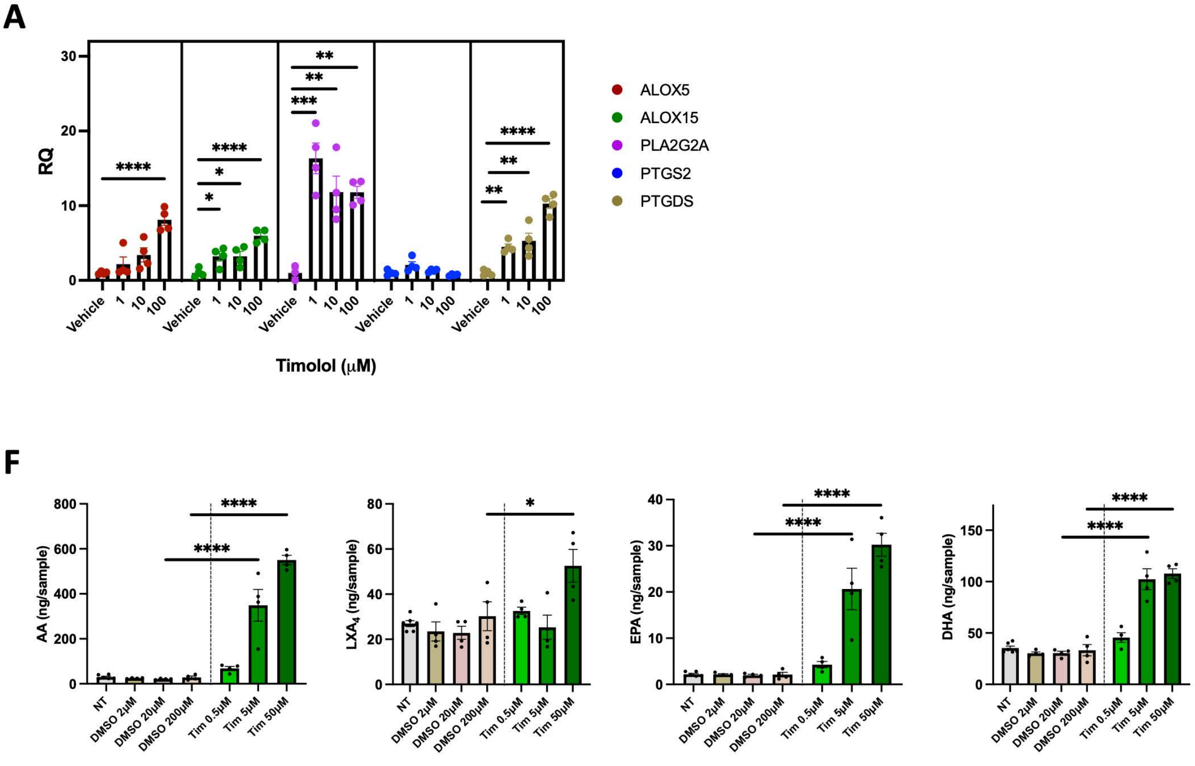


**Supplementary Figure 3. Key lipid mediator synthetic enzymes and synthesis were induced by timolol.** (**A**) Treatment with increasing concentrations of timolol significantly upregulated expression of ALOX5, ALOX15, PLA2G2A and PTGDS. (**B**) Lipidomic analyses of the timolol treated culture media showed a significant, dose dependent increase in arachidonic acid and $LXA_4$ levels, as well as EPA and DHA substrates. (*$p<0.05$, **$p<0.01$, ***$p<0.005$, ****$p<0.001$, bars are SE).

Supplementary Figure 4

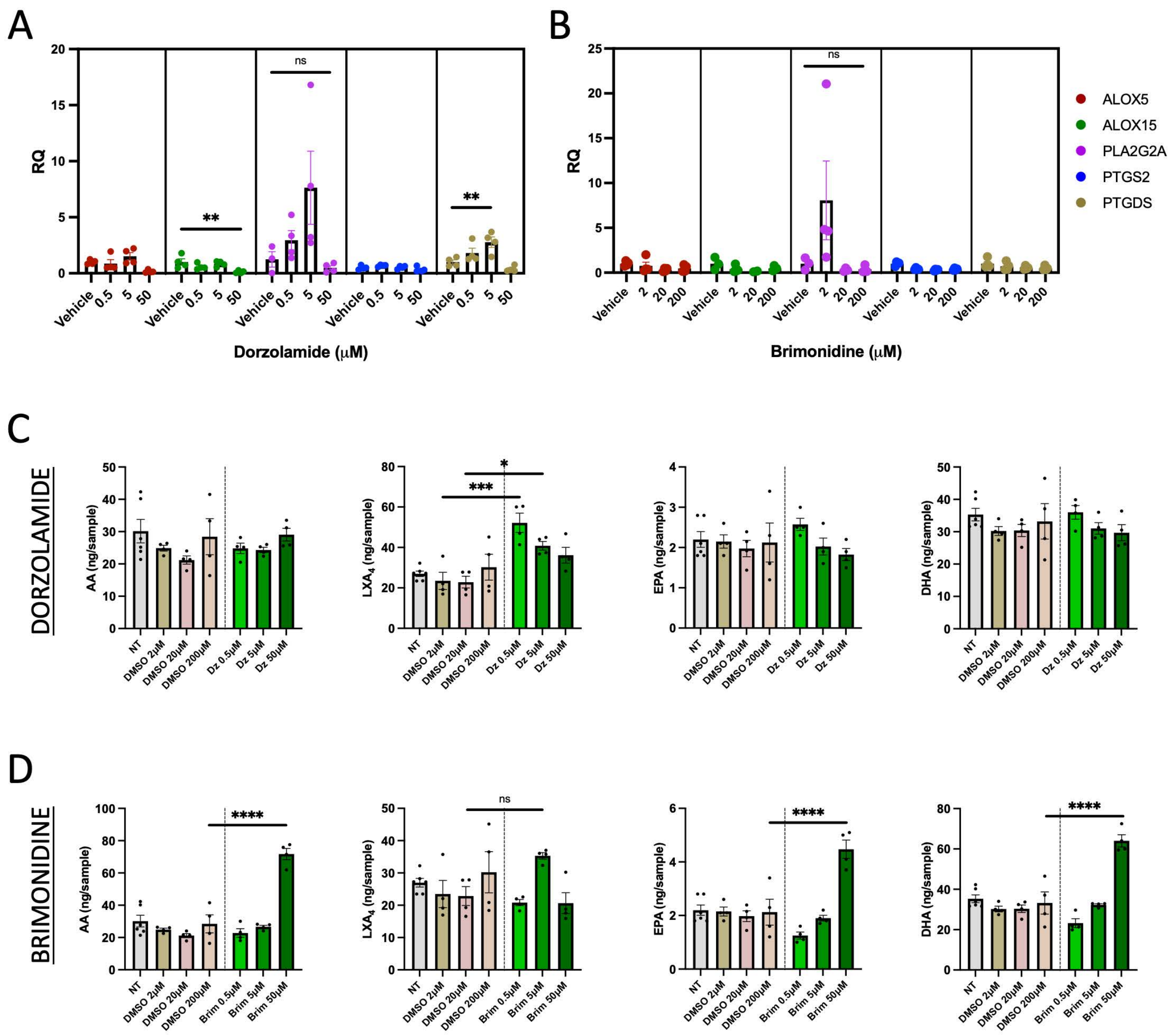


**Supplementary Figure 4. Key lipid mediator synthetic enzymes and synthesis were not strongly affected by dorzolamide or brimonidine.** (**A-B**) In contrast to Latanoprost and Timolol, Dorzolamide treatment had little effect on a panel of key synthetic enzymes, resulting in downregulation of all transcripts at 50 μm (A). Similarly, Brimonidine treatment did not result in any significant changes (B). (**C-D**) Concentrations of PUFA precursors and $LXA_4$ from analyses of TM cell culture media following treatment with dorzolamide or brimonidine. (*p<0.05, **p<0.01, ***p<0.005, ****p<0.001, bars are SE).

Supplementary Figure 5

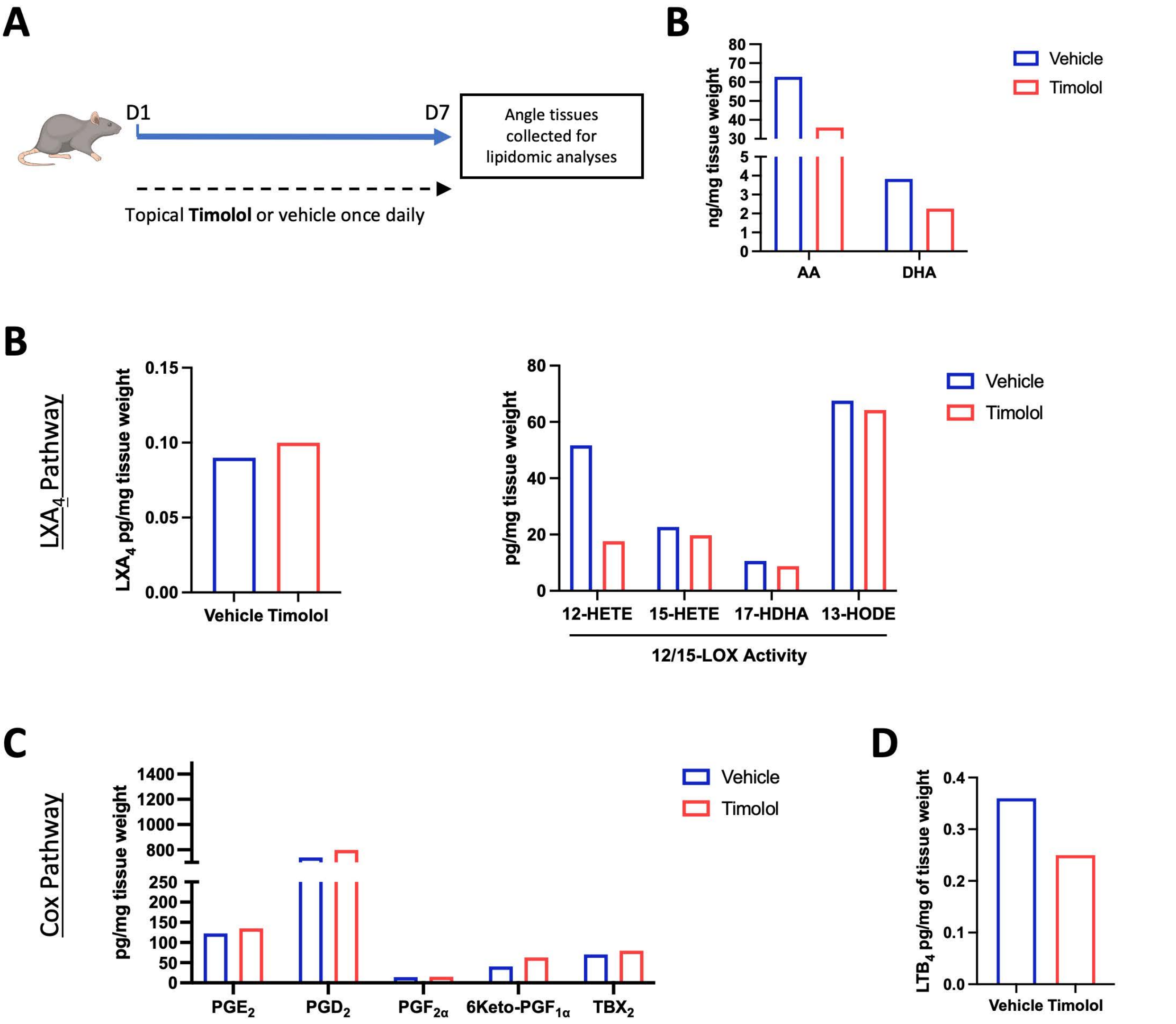


**Supplementary Figure 5. Analyses of $LXA_4$ and COX pathway mediators after treatment with timolol shows no induction *in vivo*.** A) Rats were dosed topically with timolol or vehicle daily for 7 days and angle tissues were collected and pooled for lipidomic analyses. B) Concentrations of PUFAs detected in angle tissues. C) Concentrations of products and intermediates in the $LXA_4$ pathway. D) Concentrations of Cox pathway products. E) Concentration of LTB4.
