## Supplementary Tables for "Lipidomic Analysis Reveals Drug-Induced Lipoxins in Glaucoma Treatment"

**Supplementary Table 1:** Aqueous humor lipidomic analysis results (values in pg/100 μL, *indicates a significant difference between glaucoma and control groups).

| **Analyte** | **Glaucoma** | **Control** |  | **Analyte** | **Glaucoma** | **Control** |
| --- | --- | --- | --- | --- | --- | --- |
| **AA*** | **1328.04 ± 312.43** | **643.07 ± 127.15** |  | 15-deoxy PGJ_2_ | ND | ND |
| DHA | 212.03 ± 185.33 | 131.09 ± 21.76 |  | **LXA_4_*** | **1.05 ± 0.36** | **0.74 ± 0.08** |
| EPA | 6.21 ± 5.59 | 3.76 ± 0.31 |  | LXB_4_ | ND | ND |
| 5-HETE | ND | ND |  | LTB_4_ | ND | ND |
| 12-HETE | ND | ND |  | 6-trans-LTB_4_ | ND | ND |
| 15-HETE | ND | ND |  | 20-hydroxy-LTB_4_ | ND | ND |
| 20-HETE | ND | ND |  | 20-carboxy LTB_4_ | ND | ND |
| 5-oxo-ETE | ND | ND |  | LTB_6_ | ND | ND |
| 4-HDHA | ND | ND |  | LTC_4_ | ND | ND |
| 7-HDHA | ND | ND |  | LTD_4_ | ND | ND |
| 14-HDHA | ND | ND |  | LTE_4_ | ND | ND |
| 17-HDHA | ND | ND |  | RvD_1_ | ND | ND |
| **12-HEPE*** | **1.38 ± 0.62** | **ND** |  | RvD_2_ | ND | ND |
| 15-HEPE | ND | ND |  | RvD_3_ | ND | ND |
| 18-HEPE | ND | ND |  | RvD_5_ | ND | ND |
| **13-HODE*** | **12.17 ± 3.71** | **16.32 ± 1.27** |  | RvE_1_ | ND | ND |
| PGE_2_ | 9.05 ± 12.28 | ND |  | TXB_2_ | ND | ND |
| PGD_2_ | 12.98 ± 7.77 | ND |  | NPD_1_ | ND | ND |
| PGF_2a_ | ND | ND |  | Maresin-1 | ND | ND |
| 6-keto-PGF_1a_ | ND | ND |  | Maresin-2 | ND | ND |

**Supplementary Table 2:** Rodent ocular angle tissue cytokine analysis results with and without lipoxin A_4_ treatment (*indicates a significant difference between vehicle and LXA_4_ groups).

| **Analyte** | **Vehicle (pg/mL)** | **LXA_4_ (pg/mL)** | **Analyte** | **Vehicle (pg/mL)** | **LXA_4_ (pg/mL)** |
| --- | --- | --- | --- | --- | --- |
| TGF-β_1_ | 23.16±2.27 | 27.98±2.57 | IL-6 | 330.65±30.95 | 303.18±28.80 |
| TGF-β_2_ | 123.68±11.89 | 134.07±8.44 | IL-10 | 13.34±0.84 | 12.82±1.28 |
| **TGF-β_3_*** | **9.78±1.17** | **15.79±1.98** | **IL-12p70*** | **12.60±1.68** | **5.73±1.54** |
| EGF | 7.77±1.61 | 5.99±2.06 | IL-13 | 6.90±1.21 | 5.53±0.80 |
| Eotaxin | 2.92±0.26 | 2.86±0.25 | IL-17A | 2.37±0.24 | 2.56±0.36 |
| Fractalkine | 132.57±14.13 | 125.14±11.08 | IL-18 | 3954.39±329.4 | 3452.56±209.04 |
| G-CSF | 2.07±0.20 | 2.61±0.34 | IP-10 | 11.99±0.80 | 11.82±1.32 |
| GM-CSF | 10.627±2.19 | 13±2.16 | Leptin | 257.24±35.01 | 194.41±19.57 |
| GRO/KC | 35.987±8.86 | 59.03±6.13 | LIX | 17.67±0.95 | 14.65±1.23 |
| IFN-γ | 114.15±4.83 | 104.44±6.27 | MCP-1 | 55.68±11.32 | 76.79±11.72 |
| IL-1α | 21.40±5.37 | 14.88±2.11 | **MIP-1α*** | **3.41±0.26** | **2.75±0.16** |
| IL-1β | 45.71±3.56 | 44.24±3.21 | MIP-2 | 11.25±1.88 | 11.67±3.32 |
| IL-2 | 12.64±1.69 | 9.23±1.25 | RANTES | 10.56±1.38 | 12.76±1.57 |
| IL-4 | 9.55±1.45 | 9.53±1.32 | **TNF-α*** | **7.55±0.95** | **4.12±0.60** |
| IL-5 | 22.65±1.64 | 19.91±0.97 | VEGF | 30.64±2.45 | 33.69±1.59 |
